# Computational Investigation of O_2_ Diffusion Through an Intra–molecular Tunnel in AlkB; Influence of Polarization on O_2_ Transport

**DOI:** 10.1101/154674

**Authors:** Hedieh Torabifard, G. Andrés Cisneros

**Affiliations:** Department of Chemistry, Wayne State University, Detroit, MI 48202; Department of Chemistry, University of North Texas, Denton, TX 76203

## Abstract

*E. Coli* AlkB catalyzes the direct dealkylation of various alkylated bases in damaged DNA. The diffusion of molecular Oxygen to the active site in AlkB is an essential step for the oxidative dealkylation activity. Despite detailed studies on the stepwise oxidation mechanism of AlkB, there is no conclusive picture of how O_2_ molecules reach the active site of the protein. Yu *et al. (Nature,* **439**, 879) proposed the existence of an intra–molecular tunnel based on their initial crystal structures of AlkB. We have employed computational simulations to investigate possible migration pathways inside AlkB for O_2_ molecules. Extensive molecular dynamics (MD) simulations, including explicit ligand sampling and potential of mean force (PMF) calculations, have been performed to provide a microscopic description of the O_2_ delivery pathway in AlkB. Analysis of intra–molecular tunnels using the CAVER software indicates two possible pathways for O_2_ to diffuse into the AlkB active site. Explicit ligand sampling simulations suggests that only one of these tunnels provides a viable route. The free energy path for an oxygen molecule to travel along each of these tunnels has been determined with AMBER and AMOEBA. Both PMFs indicate passive transport of O_2_ from the surface of the protein. However, the inclusion of explicit polarization shows a very large barrier for diffusion of the co–substrate out of the active site, compared with the non–polarizable potential. In addition, our results suggest that the mutation of a conserved residue along the tunnel, Y178, has dramatic effects on the dynamics of AlkB and on the transport of O_2_ along the tunnel.

## 1 Introduction

*E. Coli* AlkB is a member of the Fe(II)/2–oxoglurarate dependent enzyme superfamily. AlkB is in charge of the repair of alkylated DNA and RNA bases by catalyzing an oxidative dealkylation mechanism.^1–3^ The oxidative dealkylation mechanism catalyzed by AlkB, has been investigated extensively.^4–18^ The mechanism can be separated into two stages: The first stage involves the formation of a ferryl (Fe(IV)–oxo) intermediate, followed by the oxidation of the alkyl moiety on the damaged base. The reaction mechanism for the dealkylation of 1–methyladenine (1meA) by AlkB is shown in Scheme 1. The rate limiting step in this cycle corresponds to the abstraction of a hydrogen atom from 1meA by the ferryl moiety.^5,18^ To initiate the oxidation of Fe, an O_2_ molecule needs to displace a water molecule from the primary coordination sphere of Fe. The diffusion of O_2_ into the active site is an essential process in this mechanism. A key question in this first stage of the reaction is how O_2_ molecules are transported into the active site. Various enzymes that use molecular Oxygen have been shown to employ transient intra–molecular tunnels formed by flexible hydrophobic residues to transport O_2_ from the surface of the protein to the active site.^19–22^

**scheme 1:**
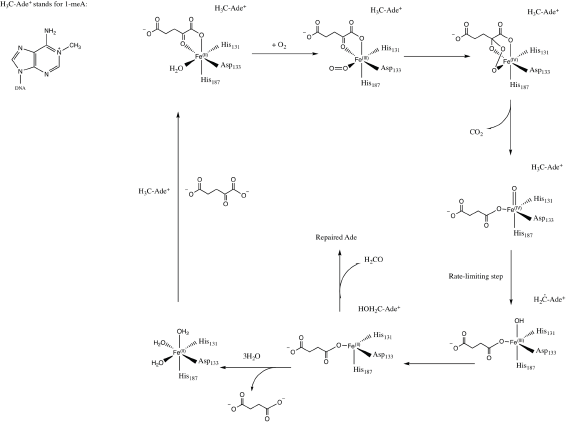
Proposed mechanism for the dealkylation of 1meA by AlkB.

A number of computational studies have been reported on the diffusion of O_2_ through intra–molecular tunnels.^22–29^ One possible approach to investigate the transport of these types of molecules involves standard MD simulation of ligand diffusion, which ideally requires a large number of independent replicate runs of several ns to attain adequate sampling (flooding simulation).^19,22^ A second approach involves the determination of the potential of mean force (PMF).^22,30^ Extensive sampling is essential to obtain accurate PMFs since incomplete sampling can result in large errors in the calculated barriers. In recent years, GPU computing has resulted in a large increase in the timescales of MD simulations, which make it possible to perform the sufficient sampling that is necessary to achieve a realistic description of ligand diffusion.^21,31^

Yu *et al.* proposed the possibility of O_2_ transport through an intra–molecular tunnel in AlkB based on the first AlkB crystal structures.^32^ They showed that there is little unoccupied volume in the binding cavity, in which the target base directly contacts its molecular surface. This study portrays open and closed states of a tunnel putatively gating O_2_ diffusion into the active site in the presence of iron(II) and cobalt(II), respectively. The open and closed states of this tunnel are due to the structural variation of the side chain of residue W178, which is located at the entrance of a binding cavity.^33^

Based on these observations, we present results from various computational approaches to determine the likelihood of the existence of this putative intra–molecular tunnel, and whether it is amenable for the passive transport of O_2_ into the active site of AlkB. In addition, given that molecular Oxygen is a neutral albeit highly polarizable molecule, we have employed both non–polarizable fixed–charge (AMBER) and multipolar/polarizable (AMOEBA) potentials to investigate the role of electronic polarization. Explicit simulations on three W178 mutants have also been performed to investigate the role of this particular residue on AlkB structure and the structure of the tunnel. The remainder of the paper is organized as follows: Section 2 presents the details for the various methods employed to simulate the transport of the substrate, calculate the energy for this transport along the tunnels, and various analysis. Subsequently, section 3 describes the results and discussion for the calculated tunnels and transport/energetics, followed by concluding remarks.

## 2 Computational Methods

The initial structure for wild type *E. Coli* AlkB in complex with DNA was taken from the Protein Data Bank (PDB ID: 2FDG^32^). Hydrogen atoms, counter ions, and TIP3P water molecules were added to the holo structure with the LEaP package^34^ from AMBER14.^35^ The final system size was ≈ 50000 total atoms with 3 counter ions. This system was initially equilibrated in our previous studies on AlkB.^5,6^ MD simulations were performed with ff99SB^36^ and AMOEBA using the NPT ensemble.^37^ Simulations involving the ff99SB potential were performed with the PMEMD.cuda program from AMBER14.^35^ AMOEBA simulations were carried out with an in–house, AMOEBA capable, development branch of AMBER based on pmemd termed pmemd.gem.

All simulations used a 1 fs step size and a 9 Å cutoff for non–bonded interactions. The non–polarizable simulations were run at 300 K using Langevin dynamics^38^ (*γ*=1 ps^-1^), with a Berendsen barostat.^39^ The parameters for 1–methyladenine (1meA), α–KG and O_2_ for both ff99SB and AMOEBA were developed in–house and are provided as Supplementary Information. The iron cation was approximated by Mg(II) parameters based on the precedent established by our previous studies on AlkB ^5,6^ and TET2.^40^ SHAKE^41^ was used for all the simulations and the smooth particle mesh Ewald (PME) method^42^ was employed to treat long–range Coulomb interactions.

The existence of possible tunnels for O_2_ transport in AlkB were determined by analyzing the crystal structure, as well as 250 snapshots out of a 50 ns simulation from the non–polarizable potential using CAVER^43^ as implemented in PyMOL.^44^ Once the coordinates of the tunnel were obtained, umbrella sampling^45^ and WHAM^46–48^ were used to calculate the potential of mean force for the transport of Oxygen along the tunnel. Two possible channels were obtained from CAVER analysis.^49^ Each of these channels was populated by positioning ≈ 30–40 oxygen molecules to achieve a spacing of 0.3-0.5 Å between adjacent umbrella windows. A harmonic potential of 10 kcal mol^-1^ Å^-2^ was applied to restrain the motion of the oxygen molecule in each window. The PMF coordinate was set as the distance between the oxygen molecule and T208 (OG1) for the blue tunnel, and *α*–KG (C2) in the red tunnel. Each window was sampled for 1 ns in explicit solvent (TIP3P water model) at 300 K in NPT ensemble using Langevin thermostat and Beredsen barostat. The free energy associated with the oxygen transport along each tunnel were calculated using 1DWHAM (one-dimensional weighted histogram analysis method) technique with bootstrapping.^46–48,50,51^ in AlkB were determined by All 1DWHAM calculations were performed with the WHAM package. ^52^

The PMF was also calculated using the multipolar-polarizable AMOEBA potential . The use of higher-order multipoles and/or explicit polarization has been shown to provide accurate description of various system such as water,^53–55^ organic molecules,^56,57^ peptides,^58^ protein–ligand binding^59^ and ion channels.^60^ Several methods have been developed to include many-body polarization, such as the fluctuating charge approach,^61,62^ the Drude oscillator model^63,64^ and atomic induced dipole methods.^65,66^ AMOEBA combines distributed atomic multipoles (up to quadrupole) and (Tholé damped) inducible atomic dipoles, which have been shown to provide an accurate representation for various systems.^37,53,57,58,67–72^

Umbrella sampling/WHAM calculations for the AMOEBA calculations were performed using the amoebabio09 potential^73^ using the same windows as for the ff99SB calculations. A harmonic potential (10 kcal mol^-1^ Å^-2^, same as for the ff99SB PMF) was for each window. Each window was sampled for 100 ps in explicit AMOEBA/GEMDM water^55^ at 300 K in the NPT ensemble using the Berendsen thermostat–barostat.

Long-range electrostatic interactions were computed employing the smooth particle mesh Ewald method^42,74,75^ with an 8.5 Å direct cutoff. All PMFs were calculated with the WHAM program from Grossfield *et al.*^52^ Bootstrap analysis was done to validate the reproducibility of the data for each free energy calculation. The PMF calculations have been performed with and without the inclusion of the water molecule coordinated to the metal cation to investigate the possible effect on the diffusion of the O_2_ molecule by the screening of the cation’s charge by this water.

Direct O_2_ diffusion was also investigated by running long MD simulations (500 ns) with the ff99SB force field (exclusively) on wild type AlkB and three mutants in the presence of O_2_ molecules. To probe for O_2_ delivery pathways in AlkB, 10 O_2_ molecules, corresponding to 0.03 M [O_2_] calculated with respect to the total simulation box volume of 520000 Å^3^, were added to the equilibrated AlkB model. This O_2_ concentration is 23–fold higher than the saturated O_2_ concentration in water at ambient condition. This high O_2_ concentration was introduced to maximize the sampling of O_2_ delivery pathway(s) in the protein within limited time scales (500 ns). Four independent simulation systems were designed for wild type in which O_2_ molecules were initially distributed randomly around protein surface in the solution. Each system was visually inspected to ensure that the added O_2_ molecules did not overlap with atoms of other molecules, and then was simulated for 500 ns in the NPT ensemble at 300 K using Langevin dynamics (*γ* = 1 ps^-1^) coupled to a Berendsen barostat. Repeated simulations were performed to improve statistics and to provide a more accurate description of the O_2_ delivery pathway.

Three–dimensional (3D) density maps representing the O_2_ occupancy profiles were calculated to identify O_2_ delivery pathways, using the Volmap tool as implemented in the VMD program.^76^ Animations made from trajectories for the WT and all mutant simulations are provided as Supplementary Information. Root mean square deviation (RMSD), root mean square fluctuation (RMSF), hydrogen bond, correlation, histogram and distance analyses were performed using the cpptraj module available in AMBER14 suite. ^35^

Energy decomposition analysis (EDA) was used to calculate non–bonded intermolecular interaction energies (Coulomb and VdW interactions) between selected residues. All EDA calculations were carried out with an in–house FORTRAN90 program.^77,78^ The average non-bonded interaction between a particular residue, W178, and every other residue is approximated by Δ*E_int_*= 〈ΔE_*i*_〉, where *i* represents an individual residue, 〈ΔE_*i*_〉 represents the non-bonded interaction (Coulomb or VdW) between residue *i* and the residue of interest, W178, and the broken brackets represent averages over the complete production ensemble obtained from the MD simulations. This analysis has been previously employed for QM/MM and MD simulations to study a number of protein systems.^5,6,30,78–80^

## 3 Results and Discussion

This section presents the results obtained from the MD simulations and analysis for the oxygen transport through AlkB and three W178 mutants. Subsection 3.1 presents the results of the CAVER analysis for the crystal structure and selected MD snapshots to investigate the prevalence of each tunnel along the trajectory. Long MD simulations on wild type and W178 mutants are discussed in Subsections 3.2 and 3.3. This is followed by the presentation of the free energy profiles obtained from the umbrella sampling and WHAM calculations in Subsection 3.4.

### 3.1 Probability of occurrence of different tunnels in wild type AlkB

An initial analysis of the original AlkB crystal structure, 2FDG,^32^ using CAVER indicates the existence of four possible tunnels (Figure 1a–Figure S1 a). All tunnels start from the surface of the protein and reach Fe(II) in the active site. The CAVER analysis further indicates that the blue and red tunnels are the most probable tunnels for O_2_ diffusion due to their high throughput, low curvature and short length (Table S1). The blue tunnel is defined by residues L15, A16, A17, A19, I143, S145, F154, F156, K166, W178, S182, F185 and H187. The red tunnel which comprises residues H131, Q132, D133, L139, I143, W178, S182, R183, L184, F185, H187, and R210 is consistent with the proposed tunnel by Yu *et al.*^32^ Figures 1b and 1c show the residues that define the red and blue tunnels. Figure S1 b shows the position of the blue and red tunnels with respect to the AlkB active site. The coordinated water molecule trans to H131 occupies the site that would be replaced by the oxygen molecule traveling along each tunnel.^81^

**Figure 1:**
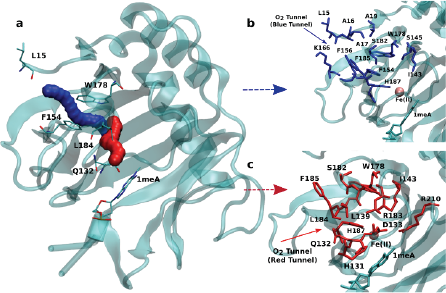
a) Two main tunnels obtained with CAVER for the crystal structure of AlkB. Residues that define the blue (b) and red (c) tunnels.

Given that the initial CAVER analysis is based on a static structure, further analysis was performed to investigate the effects of the protein motion on the predicted tunnels. To this end, a short MD simulation (50 ns) was carried out on wild type AlkB and 250 random snapshots were extracted and subjected to CAVER analysis. The results for all 250 snapshots indicate that out of these samples 29.4*%* exhibit the availability of the blue tunnel compared to 28.2*%* occurrence of the red tunnel (Table S2). In addition, the CAVER analysis suggests that the average curvature of the blue tunnel is smaller than that of the red tunnel. Taken together, these results indicate that O_2_ molecules should be more easily transported through the blue tunnel.

The CAVER algorithm simplifies the oxygen molecules and protein atoms as hard spheres whose sizes are approximated by van der Waals radii. The algorithm determines the existence of putative tunnels purely based on steric considerations, without considering actual intermolecular interactions. This steric nature of the calculation prevents any insight into the energetics of transport along the tunnel.

To further test the CAVER prediction, long MD simulations (500 ns) on wild type AlkB in the presence of 10 O_2_ molecules randomly positioned in the surface of the protein have been performed to investigate how O_2_ molecules travel along these tunnels. The results strikingly confirmed the prediction and show that the O_2_ molecules prefer to travel along the blue tunnel rather than other tunnels as described in the next Subsection.

### 3.2 Long MD simulations reveal the main oxygen diffusion pathway in AlkB

Four independent MD simulations on wild type AlkB were performed to investigate the diffusion of O_2_ into and out of the active site. Each of the four independent simulations contained 10 O_2_ molecules and extended for 0.5 *μ*s. During these simulations we observe 5 O_2_ molecules diffuse into the active site on average. In all cases the diffusion of the ligand occurs exclusively via the blue tunnel pathway (see movie S1 for animation).

Figure 2a shows the 3D density map representing the occupancy profile for O_2_ molecules for the wild type and three W178 mutants (see below for mutant simulations). This density map is an average over all four 0.5 *μ*s trajectories. The calculated density maps show that the cavities occupied by O_2_ molecules in the wild type are consistent with the coordinates of the blue tunnel. The distance between the Fe(II) atom and O_2_ molecules for each independent simulation are presented in Figure S2. Distances smaller than 6 Å between an O_2_ molecule and Fe(II) were considered as a complete entrance. In all wild type simulations we observed that all Oxygen molecules that diffuse into the active site spend a very short time close to the Fe(II) atom on average, and then they escape from active site and return to solution.

**Figure 2:**
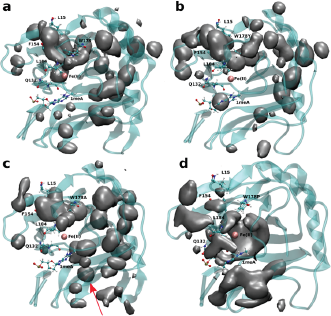
3D density map representing the O_2_ molecule occupancy in a) WT, b) W178Y, c) W178A, d) W178P. The O_2_ molecules diffuse through the blue tunnel for wild type and W178Y. W178A reveals a new pathway beside the blue tunnel indicated by a red arrow. O_2_ molecules are delivered through the blue tunnel and exhibit long residence times around Fe(II) in the W178P mutant. The isovalue for all density maps is 0.006.

### 3.3 Mutation of W178 can change the diffusion pathway

Crystal studies on AlkB^32^ show that residue W178 is located at the entrance of the binding cavity. Based on its location, Yu *et al.* suggested that Y178 could act as a gate along the tunnel depending on its various side chain conformations. The effect of W178 on O_2_ diffusion was probed by performing MD simulations on three AlkB mutants: W178Y, W178A and W178P. These simulations were subjected to various analyses including density maps, RMSD, RMSF and energy decomposition to investigate the effect of the mutation on the structure and dynamics of AlkB.

The occupancy profiles for each mutant are presented in Figures 2b–d. The occupancy profile reveals that O_2_ molecules in W178Y mutant diffuse through the blue tunnel similarly to wild type. The correlation analysis by residue (Figure S5 e) indicates that W178Y has a pattern similar to that of the wild type, suggesting that the dynamics of this mutant and the wild type AlkB are similar. However, distance analysis (Figure S3) indicates that O_2_ molecules in the W178Y mutant spend more time in the active site compared to the wild type (Figure S2). This difference in residence time could be due to the difference in size and flexibility of the tyrosine side chain with respect to tryptophan. All systems are observed to be stable throughout the simulation time as per RMSD with respect to the crystal structure (Figure S4).

The analysis of the change in fluctuation along the calculated trajectories between the wild type and each mutant shows striking differences. Figure 3a suggests that residues around the blue tunnel in the W178Y mutant fluctuate less than in the wild type. Reduced fluctuation of these residues may explain why O_2_ molecules spend more time in the active site compared to the wild type.

**Figure 3:**
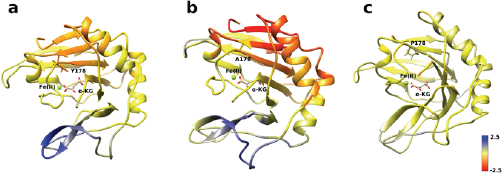
RMSF difference between wild type and mutants along the sampled trajectories for (a) W178Y, (b) W178A and (c) W178P. Residues in blue(red) denote larger(smaller) fluctuations in the mutant structures compared to the wild type.

Similar to the wild type and W178Y mutant, the O_2_ diffusion in the W178P mutant occurs only through the blue tunnel. However, the occupancy profile (Figure 2d), distance analysis (Figure S3) and the movie of the trajectories (movie S3) show that O_2_ molecules spend a significant portion of the simulation time around the Fe(II) atom in the active site, indicating that O_2_ gets trapped in the active site for a larger period. The RMSF (Figure 3c) shows that the residue at the 178 position exhibits larger fluctuations in the P mutant than in the wild type structure.

The main pathway for oxygen diffusion in the W178A mutant is the blue tunnel. However, this mutant reveals a new pathway to transport O_2_ molecules from the surface of the protein into the active site (Figure 2c). The correlation difference plot (Figure S5 g) shows that residues 163 to 189 in the W178A mutant and wild type are anti–correlated. In addition, the RMSF difference analysis (Figure 3b) shows increased fluctuation for the residues around the new pathway in the W178A mutant compared to the wild type. The increased fluctuation of these residues may help explain for appearance of a new pathway in the W178A mutant.

Energy decomposition analysis (EDA) was carried out to further understand residue–by–residue inter-molecular interactions for the wild type and the three mutant systems. The EDA analysis suggests that the mutation of tryptophan to tyrosine results in a negligible change to the stability of this residue with respect to the rest of the the protein (Table 1), however, the stability of the whole protein increases significantly (Table S3). In the case of the W178P system, the EDA analysis reveals less stability for site 178 when P occupies this site in this mutant compared to wild type (Table 1). The sum of all intermolecular interactions in this mutant compared to wild type (Table S3) suggests that the mutation of tryptophane to proline results in an overall destabilization of the protein.

**Table 1.** Intermolecular energy difference analysis between residue 178 and all protein and DNA residues for the wild type and W178Y/A/P mutants. The values for the wild type correspond to the average over all 4 independent simulations (all energies in kcal/mol).

| | $\Delta E_{Coul}$ | $\Delta E_{vdW}$ | $\Delta E_{Tot}$ |
| --- | --- | --- | --- |
| WT | $-41.2 \pm 0.8$ | $-23.7 \pm 0.4$ | $-64.9 \pm 0.6$ |
| W178Y | -43.7 | -21.4 | -65.1 |
| W178A | -41.8 | -9.7 | -51.5 |
| W178P | -32.3 | -13.8 | -46.1 |

In summary, the MD simulations indicate that the main pathway for oxygen diffusion in all mutants is the blue tunnel. However, the access to this tunnel can be facilitated or hindered by the mutation of W178, depending on the size and flexibility of the residue and the resulting effect of the mutation at this position on the structure and dynamics of AlkB. These results support the importance of W178 for oxygen diffusion, and provide an interesting site to test experimentally in the future.

### 3.4 PMF calculation for O_2_ pathway to the active site

As described in Subsection 3.2, the diffusion MD simulations suggest that the O_2_ molecules diffuse to the active site through the blue tunnel exclusively, and remain in the active site for a short time before returning to solution. Therefore, we have calculated the PMFs associated with passive O_2_ transport through both tunnels in order to gain further insights for the possible reason for the tunnel preference and causes for the short residence times of O_2_ molecules in the active site.

The calculated PMFs for the transport of molecular oxygen along the blue and red tunnels (the coordinated water is included in the calculations) using the ff99SB potential are shown in Figure 4 (see Figure S6 and S7 for histogram analysis, and Figures S9 and S10 for bootstrap analysis). The PMF results suggest that transport of Oxygen molecules into the active site is downhill by ≈ 3 kcal/mol through the blue tunnel and the Oxygen molecules only need to overcome a small free energy barrier (less than 1 kcal/mol) around 6-7 Å from the active site and . Conversely, O_2_ that diffuses through the red tunnel need to overcome a barrier of ≈ 1.5 kcal/mol to reach the active site at the entrance of the tunnel.

**Figure 4:**
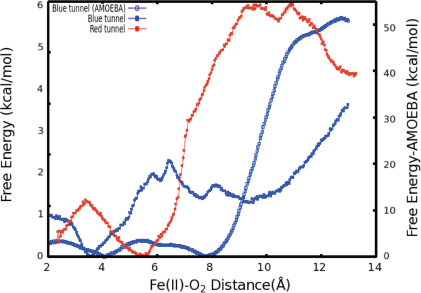
Calculated PMFs for intramolecular O_2_ transport in AlkB. The PMF for the blue tunnel was calculated with the ff99SB, and AMOEBA force fields. For the PMF associated with transport along the red tunnel only the ff99SB potential was used. The coordinated water is included in this calculations.

The barrier observed along the red tunnel likely corresponds to the interaction between some residues with each other along this tunnel. For instance three residues along the red tunnel, E136, R183, and R210, form hydrogen bonds for more than 30% of the simulation time (Table S4). These residues are located around 9–11 Å from the active site and correspond to the region where the O_2_ molecules experience the free energy barrier. The interaction between this residues could make the O_2_ diffusion more difficult as the co–substrate molecules need to break these inter-residue hydrogen bonds in order to transit through this region of the red tunnel. The calculated barrier in the red tunnel compared to a completely downhill energy path in the blue tunnel provides a possible explanation for the preference of O_2_ molecule diffusion through the blue tunnel rather than the red tunnel. These results are also consistent with the long MD simulations, where O_2_ diffusion for wild type AlkB was observed only along the blue tunnel.

Furthermore, the relatively small barrier in the PMF for the blue tunnel for the egress of O_2_ from the active site provides a possible explanation as to why oxygen molecules are observed to easily escape from the active site and go back to the solution using the ff99SB potential in the free diffusion calculations (see movie S1). However, if the binding energy for O_2_ is too small it can be expected that this would drive the equilibrium toward the first stage of the reaction (scheme 1), possibly making the binding of O_2_ the rate limiting step. Thus, the short residence time of O_2_ molecules in the active site is inconsistent with the available experimental and computational results on the mechanism of AlkB, since it is known that the rate limiting step is the oxidation of the alkyl moiety by the ferryl intermediate.^5,18^

Therefore, a question arises regarding the accuracy of the PMF based on the nonpolarizable potential. In particular, whether this force field can provide a sufficiently accurate description of the inter-molecular interactions given the fact that O_2_ is a neutral molecule and therefore most fixed–charge potentials can only represent it by Van der Waals interactions. Moreover, it is known that molecular Oxygen is highly polarizable. Therefore, we performed PMF calculations on the main proposed pathway (blue tunnel) with the multipolar–polarizable AMOEBA potential to examine the effect of polarization on a highly polarizable system.

When explicit polarization is taken into account, the reduction in free energy from the surface of the protein to the active site is calculated to be ≈ 51.5 kcal/mol as shown in Figure 4 (see Figure S8 for histogram analysis and Figure S11 for bootstrap analysis). The PMFs without the coordinated water in the primary coordination sphere are presented in Figure S12. The histogram and bootstrapping analysis for all PMFs are presented in Figures S13–S18. Overall, the shape of the PMF is similar in both systems (with or without a coordinated water to the metal).

By contrast with the results obtained with the ff99SB force field, this large free energy change along the tunnel indicates that once O_2_ molecules diffuse into the active site, it more than likely stays inside the protein and binds to Fe(II) to drive the oxidation forward. This large free energy barrier precludes the egress of molecular Oxygen from the active site, which enhances the likelyhood of O_2_ molecules binding to the Fe(II) cation in the active site. The function of AlkB (and ALKBH homologues) should also be considered. Indeed, these enzymes are adaptive–response proteins since they are only expressed in response to a particular event, DNA alkylation damage, and are not present in high concentration in normal tissues under non–stress conditions. Moreover, these enzymes require a second co–factor (*α*–kg) and a substrate (1mA for AlkB) to turn over, and are mainly located in the nuclei (and to a lesser degree in mytochondria) where the oxygen concentration is known to be low.^82^ Thus, a high affinity for the co-substrate would be beneficial. Although we are not able to perform ligand diffusion simulations based on long MD calculations using the AMOEBA force field due to computational constraints, our results underscore the importance of the inclusion of polarization in certain biological systems.

## 4 Conclusion

Various computational approaches have been employed to gain insights on the transport of molecular Oxygen through intra–molecular tunnels in AlkB. Tunnel analysis based on steric considerations indicates the existence of two possible tunnels in AlkB. Ligand diffusion simulations based on long MD trajectories using a non–polarizable fixed–charge potential (ff99SB) show that only one of these pathways is observed during the simulations on the wild type enzyme. O_2_ diffusion in the W178Y/P mutants takes place through the same main tunnel (blue tunnel), while simulations on the W178A mutant reveal a new pathway. The modeling shows that the replacement of W178 with tyrosine has only a modest effect on the O_2_ diffusion, while replacement by proline creates a barrier and O_2_ molecules can not readily bypass it. The calculated PMFs for the wild type are consistent with the diffusion simulations and show that the free energy associated with O_2_ transport along the blue tunnel is completely downhill, compared to a barrier of ≈ 3 kcal/mol for the red tunnel. However, the small barrier for O_2_ egress from the active site through the blue tunnel is inconsistent with experimental and computational results for the AlkB mechanism. Complementary PMF simulations with the polarizable AMOEBA potential indicate that the free energy of O_2_ transport when electronic polarization is taken into account remains mostly downhill, but the barrier for egress is significantly increased by almost one order of magnitude. Our results provide support for the existence of an intra–molecular O_2_ tunnel in AlkB, the role of a key conserved residue (Y178), and the importance of accounting for polarization for O_2_ transport simulations.

## Acknowledgements

This work was supported by grants R01GM108583, and R01GM118501 to GAC and by Wayne State University. Computing time from Wayne State’s C&IT and the University of North Texas (CASCaM, NSF CHE-1531468) are gratefully acknowledged.

## Supporting information

Tunnel analysis for crystal structure and MD simulations, EDA, H-bonding, distance, RMSD, correlation, histogram, bootstrapping analyses, and additional file with AMBER and AMOEBA parameters for 1meA, O_2_, and *α*-KG are provided in the supporting information.

